# Experimental galactose-1 phosphate uridylyltransferase (GALT) mRNA therapy improves motor-related phenotypes in a mouse model of Classic Galactosemia

**DOI:** 10.1101/2025.04.21.649843

**Authors:** Olivia Bellagamba, Aaron Guo, Xinhua Yan, Joe Sarkis, Bijina Balakrishnan, Kent Lai

## Abstract

Despite life-saving newborn screening programs and a life-long galactose-restricted diet, many patients with Classic Galactosemia continue to develop long-term debilitating neurological deficits, speech dyspraxia, and primary ovarian insufficiency (POI). Earlier, we showed that administration of an experimental human *GALT* mRNA predominantly expressed in the liver of the *GalT* gene-trapped mouse model augmented the expression of hepatic GALT activity, which reduced build-up of galactose and its toxic metabolites not only in the liver, but also in the peripheral tissues. Moreover, we showed that administration of *GALT* mRNA in the mutant mice restored whole-body galactose oxidation (WBGO), a functional biomarker. In this study, we extended our proof-of-concept (POC) efficacy studies to a disease-relevant phenotype, motor impairment. We found that a biweekly dosing regimen at 2mg/kg for 2 months could improve the motor performance of the animals in Rotarod and Composite Phenotype Scoring tests.

## 1. Introduction

Classic Galactosemia (OMIM 230400) is an autosomal recessive disorder caused by deficiency of <u>gal</u>actose-1-phosphate uridylyltransferase (**GALT**, EC 2.7.7.12) activity. ^1–5^ GALT is the second enzyme in the evolutionarily conserved Leloir pathway of galactose metabolism, which catalyzes the simultaneous conversion of uridine diphosphoglucose (UDP-glucose) and galactose-1 phosphate (gal-1P) to uridine diphosphogalactose (UDP-galactose) and glucose-1 phosphate.^6^ GALT deficiency leads to accumulation of toxic galactose metabolites such as gal-1P and deficiency of UDP-galactose in patient cells.^7, 8^ If undiagnosed and untreated, Classic Galactosemia can be lethal for the affected newborns.^3^ Since the inclusion of this disease in the newborn screening panel in the U.S., neonatal mortality has decreased.^9^ The mainstay of treatment is the withdrawal of galactose from the diet.^1^ However, despite early and adequate dietary management, endogenous production of galactose and its toxic metabolites continues ^10, 11^ and patients have long-term complications such as intellectual deficits in 6 year-olds or older (45% of all patients); speech delay in 3 year-olds or older (56% of all patients), motor functions deficits (tremors & cerebellar ataxia) in 5 year-olds or older (18% of all patients), and primary ovarian insufficiency (POI) (91% of all female patients).^12, 13^ Significantly reduced bone mineralization is increasingly seen in pre-pubertal patients of both sexes.^14–16^ Except for the ovarian phenotype, which affects more than 90% of the female patients, there is considerable variability among other long-term complications. To-date, there is no effective treatment available to prevent/alleviate any of the above-mentioned long-term complications. Yet, regardless of the pathophysiological mechanisms, no one will argue that the root cause of the disease is the absence of GALT enzyme activity in the patient cells. Therefore, therapeutic strategies that aim to restore the GALT enzyme activity in the patients represent a rational and direct approach to address the unmet medical needs of the patients. Among them, experimental *GALT* mRNA therapy has emerged as a promising modality.^17–19^ Indeed, we demonstrated significant efficacy of *GALT* mRNA in normalizing the disease-relevant biomarkers and restoring whole-body galactose oxidation in a mouse model of Classic Galactosemia. In this short-term study, we expanded our preclinical proof-of-concept studies to include disease-relevant phenotype – motor impairment.

## 2. Materials and Methods

### 2.1. mRNA and LNP synthesis and formulation

mRNA was synthesized and formulated in LNPs as described previously. The same biodegradable, ionizable LNP described in our previous studies was used in the current set of studies.^18^ Briefly, sequence-optimized mRNA encoding *GALT* was synthesized *in vitro* using optimized T7 RNA polymerase-mediated transcription reaction with complete replacement of uridine by N1-methyl-pseudoridine. The linearized DNA template contains the 5′ and 3′ untranslated regions (UTRs), open reading frame (ORF), and the poly-A tail. Inverted deoxythymidine (idT) were appended to the 3’ terminus of fully synthesized mRNA by phosphodiester linkage of a modifying oligo with 3’idT. The mRNA was produced with cap1 to improve translation efficiency. Free and end-stabilized mRNA were purified, buffer exchanged and sterile filtered. The mRNAs were tested for purity and capping efficacy and were found to be > 70% and > 90%, respectively. All formulations were tested for particle size, RNA encapsulation, and endotoxin and were found to be <100 nm in size, with >80% encapsulation, and <10 EU/mL endotoxin.

### 2.2. Animal model

All animal protocols and procedures were approved and conducted in full compliance with the guidelines outlined in the Guide for the Care and Use of Laboratory Animals and were approved by the University of Utah Institutional Animal Care and Use Committee (IACUC). GalT-deficient (*GalT*-KO) mice used in this study were constructed as previously described [20] and were not challenged with galactose since weaning. Mice were housed in standard laboratory cages within temperature (22°C–23°C) and humidity (30%–70%) controlled rooms under a 12:12 light: dark cycle. All mice were confirmed by molecular genotyping using previously published protocol.^20^ Both male and female animals were used for the present study and were uniformly distributed among each experiment groups.

#### *In vivo GALT* mRNA administration

*GalT*-KO mice aged 3 and 6 weeks were administered 100 µL of sterile lipid nanoparticles (LNPs) containing *GALT* mRNA *via* intravenous injection. The LNPs encapsulated mRNA were supplied by Moderna, Inc. (Cambridge, MA, USA). Each experimental group consisted of 8 mice (4 males and 4 females), which were randomly enrolled into treatment groups at time of weaning. The mice received five bi-weekly doses of *GALT* mRNA at a concentration of 2 mg/kg. No placebo or vehicle doses were administered to the WT Control or untreated *GalT*-KO groups for this study.

### 2.3. Rotarod Test

Mice underwent Rotarod testing 3- and 9-weeks following the dosing regimen. The 6-lane Rotarod and accompanying software used in this study is purchased from Maze Engineers (Skokie, IL, U.S.A.). Mice first completed a training session on the rotarod where they would walk at a constant speed of 4RPM for at least 180 seconds before starting the first trial. Three trials were conducted where the rotarod would increase from 4RPM to 40RPM at a constant acceleration of 7.2RPM^2^. Maximum speed would be reached at 300 seconds, which was also the cutoff time for each trial. Mice would remain on the stationary rotarod between trials, where they were allowed a 5-minute rest period before beginning the next. The trial would automatically end and the latency of the mouse record when they were detected to have fallen off the rod by the IR floor sensors. Test investigators would manually end the trial when a mouse was observed to complete two passive rotations, classified by the mouse holding onto the rod as it makes a full rotation without walking. To combat small sample sizes, the latency from all three trials were used to compare mRNA-treated *GalT*-KO results to WT control and untreated *GalT*-KO performances. The WT control median latency was used to categorize “under-performing” and “over-performing” animals within each treatment group. Chi-Squared analysis was then utilized to compare the performance distribution, or the number of animals in each group to fall into these two categories. Statistical significance is achieved when the number of over- and under-performing animals in either *GalT*-KO group does not follow the expected distribution demonstrated by WT control animals. Results from chi-squared tests were verified using generalized linear models and post-HOC assessments run in R software.

### 2.4. Composite Phenotype Scoring Test

The composite phenotype scoring test was conducted in accordance with the protocol demonstrated by Guyenet *et al..*^21^ This assessment consists of four parts – the ledge test, hindlimb clasping, kyphosis, and gait tests – each measured on a 0-3 scale. The overall composite score is the combined scores from each of the four assessments, where a higher score signifies a more severe manifestation of disease phenotype (score 0-12). Two blinded investigators were assigned to evaluate the composite score test where they would average the two scores given if they differed. Half-scores were granted when the behavior did not align with the full-digit score.

The ledge test requires only the home cage of the mouse and is a simple assessment of motor coordination. Each mouse was gently placed on the ledge of the cage so that its hindlimbs can grasp onto the surface. If the mouse has no obvious issues with balance and can fully support its body while walking along the ledge, it receives a score of 0. If the mouse is observed to have several missteps as it walks on the ledge but is otherwise able to maintain its balance, it receives a score of 1. A score of 2 is granted if the hindlimbs are unable to support the body and instead the mouse is dragging along the ledge using only its front limbs. Also, if the mouse has a rough landing and/or falls onto its head instead of its front paws while climbing down into the cage, it will receive a 2. If it is incapable of walking along the ledge despite all encouragement or continues to fall off without taking a full step, this will be scored a 3.

The hindlimb clasping test is conducted by lifting the mouse from the base of its tail approximately 5-10 centimeters off the ground and observing the position of the hind legs. If the hindlimbs remain splayed outward from the abdomen, it will be scored a 0. A score of 1 is classified by the mouse retracting one of its legs in towards the abdomen for more than 50% of suspension time. If both hindlimbs are partially retracted towards the abdomen, it will be given a 2. Lastly, if both legs are completely retracted and/or touching the abdomen, this will be a score of 3.

The gait test serves to evaluate motor function and coordination. Mice are simply placed on the flat surface of a fume hood and observed by investigators. If an animal fully supports itself and evenly distributes its weight while walking, it will receive a score of 0. If there is a slight limp or abnormal stride noted, a score of 1 is granted. If the mouse is seen to have its pelvis tilted inward while walking, or demonstrates “duck feet”, it has a score of 2. The most severe score is granted when the pelvis is severely tilted inward causing the animal to drag along the surface while walking, or a combination of less severe abnormalities (i.e. duck feet, limp, uneven weight distribution).

Kyphosis is classified as the loss of spinal muscle tone and is a common symptom of neurodegeneration. Mice are observed in both a sitting and walking position to assess kyphosis. If mild or no kyphosis is observed while sitting and the spine is able to fully straighten as it walks, the mouse will receive a 0 score. If mild kyphosis is exhibited while sitting but the mouse is still able to straighten its spine while walking, it will be scored a 1. If the mouse maintains mild kyphosis as it walks it will be given a 2, and if severe kyphosis is observed while sitting and walking, it scores a 3. R software was utilized to assess any changes in composite scoring behavior between treatment groups.

### 2.5. Statistical analysis

Data are expressed as means ± SEM. Microsoft Excel, GraphPad Prism 9, and R 4.4.2 software were used to analyze data. Chi-Squared analysis was utilized for comparisons of Rotarod performances, and Mann-Whitney U test was used for Composite Phenotype Scoring analysis between groups. R software was used to corroborate significant findings from these analyses.

## 3. Results

In this pilot study, we aimed to address three questions:

Will experimental *GALT* mRNA therapy improve motor-related phenotypes in a mouse model of Classic Galactosemia?

Will the improvement, if any, of motor-related phenotypes seen in the treated animals sustain after the cessation of the experimental *GALT* mRNA treatment?

Does it matter if the GALT mRNA is offered to a younger *versus* older mouse?

**Fig. 1** below showed the overall experimental design and dosing regimen that were used to answer the above questions.

**Figure 1.**
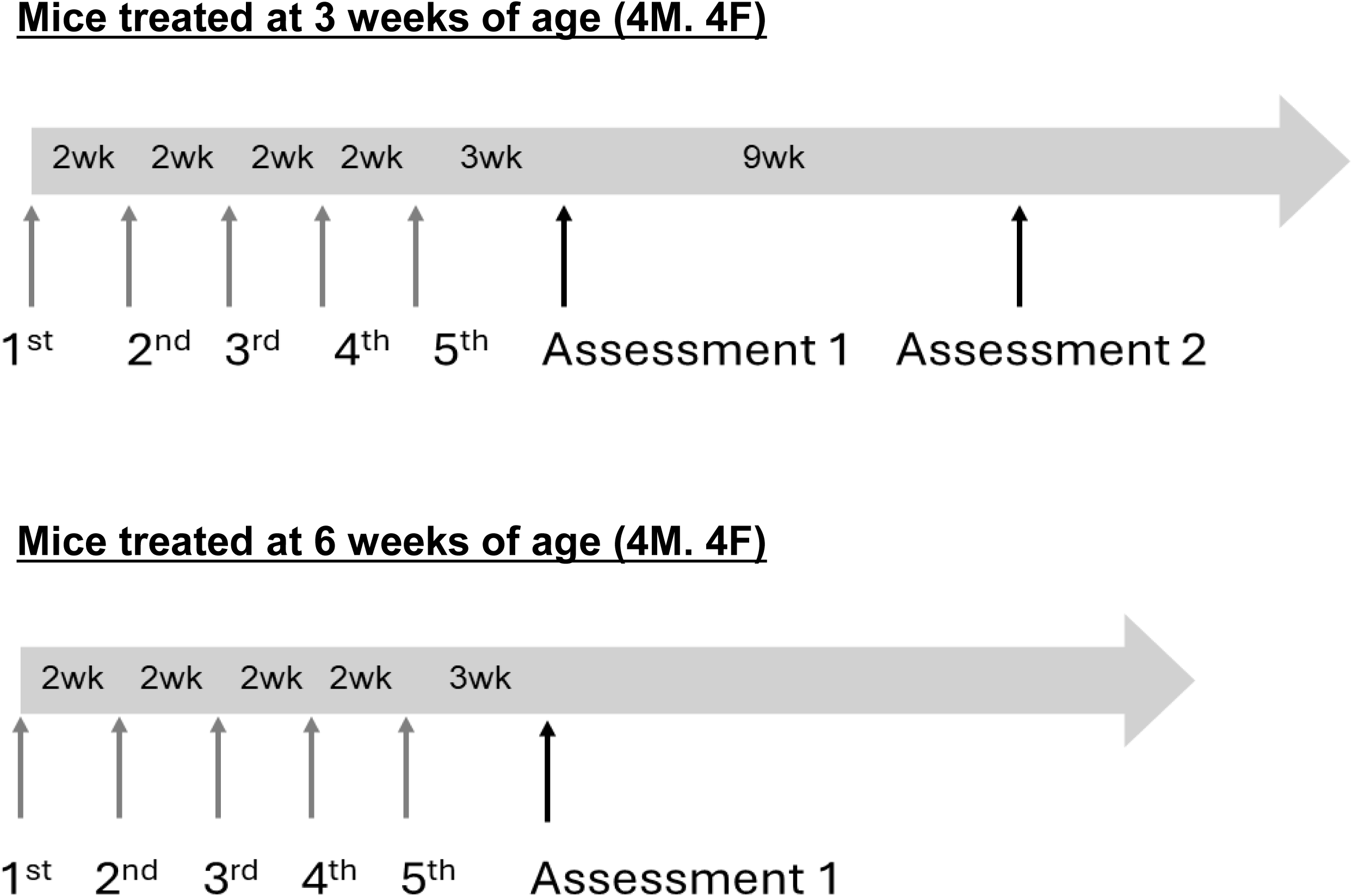
Overall experimental design for *GALT* mRNA treatment in *GalT*-KO mice. Two cohorts of *GalT*-KO mice (n=8 per cohort; 4 males, 4 females) received five bi-weekly doses (2 mg/kg) of *GALT* mRNA. Treatment began at 3 weeks of age for Cohort 1 and 6 weeks of age for Cohort 2. Rotarod and Composite Phenotype Scoring tests were conducted 3- and 9-weeks after the final mRNA dose for Cohort 1, and 3 weeks for Cohort 2 to assess motor function.

### Near-term improvement in motor functions of *GALT* mRNA treated *GalT*-KO mice

To address question **(1)**, we administered five bi-weekly doses (2mg/kg) of *GALT* mRNA to a group of *GalT*-KO mice (4 males and 4 females) starting at three weeks of age. After the last dose at 11 weeks of age, we waited for an additional 3 weeks before we performed the Rotarod and Composite Phenotype Scoring tests. Results were compared to age-matched wild type (WT) and untreated *GalT*-KO animals to determine treatment efficacy. Upon Rotarod testing, untreated WT control mice have a median latency of 106.2 seconds (**Table 1a**). WT control median latency was used to define the performances of other groups, where any treated or untreated *GalT*-KO mouse with a latency less than the WT median was labeled as “under-performing”. We then compared the number of animals that were categorized as “under-performing” and “over-performing” across treatment groups using Chi-Squared analysis. In Assessment 1, 85.7% of untreated *GalT*-KO mice performed under the WT median latency, compared to only 25% of mRNA-treated *GalT*-KO animals (**Fig 2a**). When we examined the distribution of untreated and mRNA-treated *GalT*-KO animals based on the WT control median, we see a significant improvement in the treated animals (p=0.000046). Untreated mutant mice showed a drastically different distribution compared to the WT control animals (p=0.00106). Treatment of the *GalT*-KO animals proved to be effective in diminishing the genotypic difference; in fact, we see the majority (75%) of treated *GalT*-KO mice performing over the WT median latency, which we found to be statistically significant (p=0.014) (**Fig 2a**). A generalized linear model (GLM) with a Gaussian distribution was run using treated *GalT*-KO as the reference group to confirm results obtained by Chi-Squared Testing. This model confirmed our previous findings by showing that untreated *GalT*-KO animals are estimated to perform 49.37 seconds worse than the treated *GalT*-KO group (p=0.000179) (**Supplemental Fig 1**). Post-HOC pairwise comparison further validated these results with the untreated *GalT*-KO animals having a significantly lower latency than both WT control and treated *GalT*-KO groups (p=0.0005 and p=0.0131, respectively) **(Supplemental Table 1a, b)**. Bodyweight and sex were not found to be influential factors on latency in this assessment.

**Figure 2.**
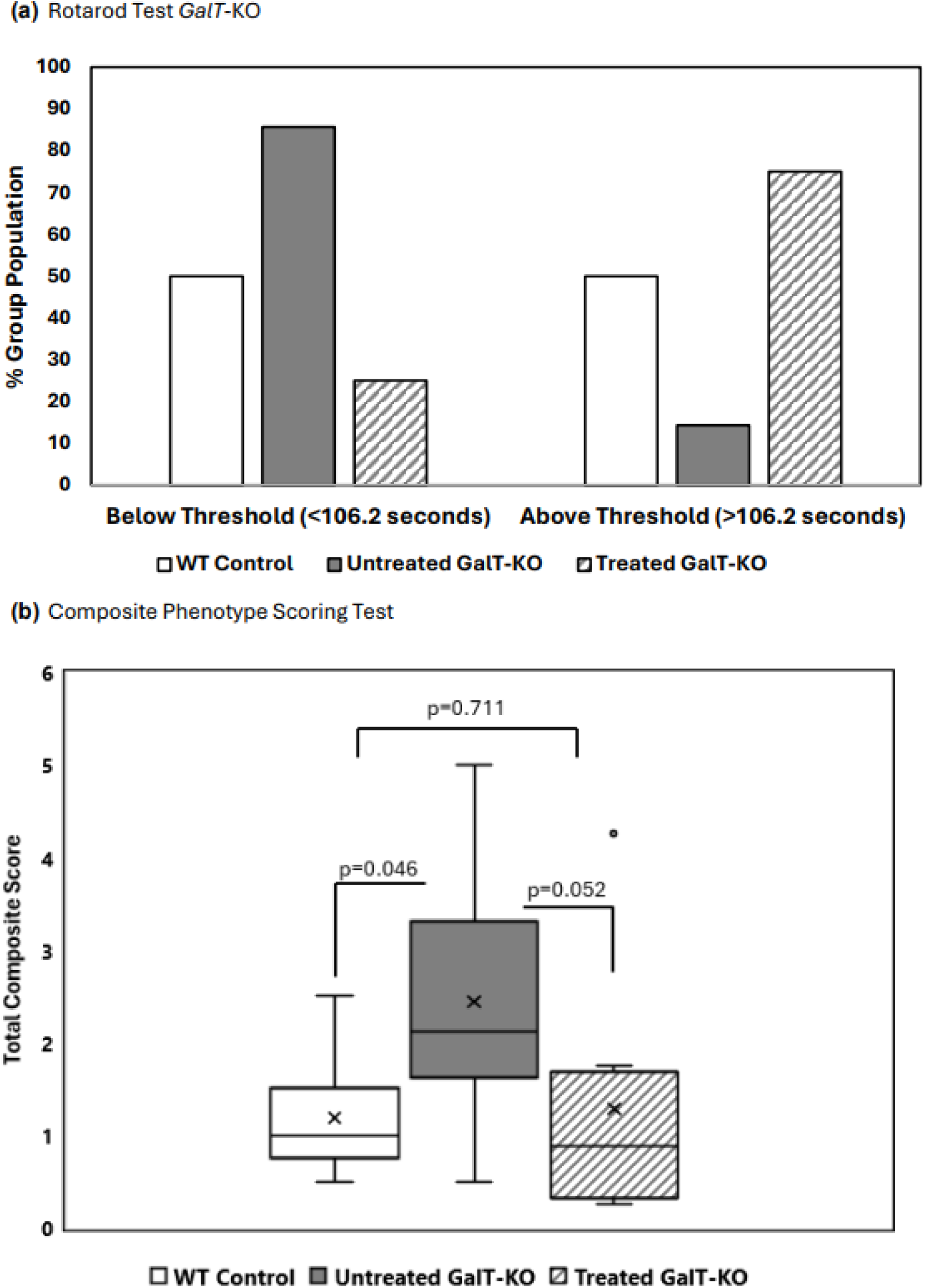
Motor function evaluation of Cohort 1 at 3 weeks after the last dose of GALT mRNA treatment. (a) Rotarod performance distribution 3 weeks after the end of treatment. The graph shows the performance distribution of each treatment group relative to the WT control group median latency. Statistical Analysis was performed using Chi-Squared Testing. (b) Composite Phenotype Scoring 3 weeks after treatment ended. The graph displays the total composite scores for each group, reflecting overall motor function and behavior. Data are presented as means ± standard error of the mean (SEM). Statistical analyses were performed using Mann-Whitney U Test. Influence of treatment group, gender, and other variables on Rotarod and Composite Score performances were analyzed by running generalized linear models with Gaussian distribution in R programming (See Supplemental Data).

**Table 1a.**
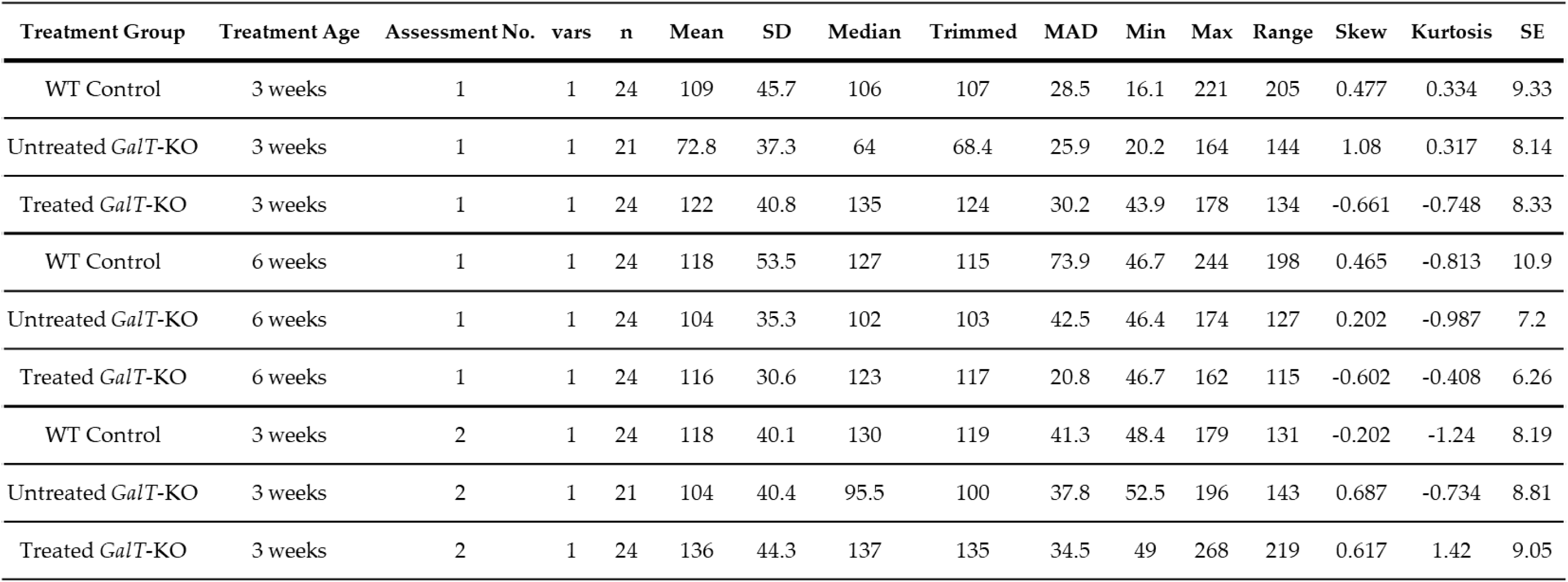
Descriptive Statistics from Rotarod Assessments.

| Treatment Group | Treatment Age | Assessment No. | vars | n | Mean | SD | Median | Trimmed | MAD | Min | Max | Range | Skew | Kurtosis | SE |
| --- | --- | --- | --- | --- | --- | --- | --- | --- | --- | --- | --- | --- | --- | --- | --- |
| WT Control | 3 weeks | 1 | 1 | 24 | 109 | 45.7 | 106 | 107 | 28.5 | 16.1 | 221 | 205 | 0.477 | 0.334 | 9.33 |
| Untreated *GalT*-KO | 3 weeks | 1 | 1 | 21 | 72.8 | 37.3 | 64 | 68.4 | 25.9 | 20.2 | 164 | 144 | 1.08 | 0.317 | 8.14 |
| Treated *GalT*-KO | 3 weeks | 1 | 1 | 24 | 122 | 40.8 | 135 | 124 | 30.2 | 43.9 | 178 | 134 | -0.661 | -0.748 | 8.33 |
| WT Control | 6 weeks | 1 | 1 | 24 | 118 | 53.5 | 127 | 115 | 73.9 | 46.7 | 244 | 198 | 0.465 | -0.813 | 10.9 |
| Untreated *GalT*-KO | 6 weeks | 1 | 1 | 24 | 104 | 35.3 | 102 | 103 | 42.5 | 46.4 | 174 | 127 | 0.202 | -0.987 | 7.2 |
| Treated *GalT*-KO | 6 weeks | 1 | 1 | 24 | 116 | 30.6 | 123 | 117 | 20.8 | 46.7 | 162 | 115 | -0.602 | -0.408 | 6.26 |
| WT Control | 3 weeks | 2 | 1 | 24 | 118 | 40.1 | 130 | 119 | 41.3 | 48.4 | 179 | 131 | -0.202 | -1.24 | 8.19 |
| Untreated *GalT*-KO | 3 weeks | 2 | 1 | 21 | 104 | 40.4 | 95.5 | 100 | 37.8 | 52.5 | 196 | 143 | 0.687 | -0.734 | 8.81 |
| Treated *GalT*-KO | 3 weeks | 2 | 1 | 24 | 136 | 44.3 | 137 | 135 | 34.5 | 49 | 268 | 219 | 0.617 | 1.42 | 9.05 |

We continued to see significant distinctions between the performances of the untreated WT and *GalT*-KO during the Composite Phenotype Scoring Test. Untreated WT control mice have a median combined score of 1, and untreated *GalT*-KO mice scored significantly higher (more disease phenotype) with a median score of 2.13 (p=0.0455) (**Table 1b**). mRNA treated *GalT*-KO mice showed improvement and scored substantially lower than the untreated *GalT*-KO cohort with a median score of 0.88 (p=0.05) (**Fig. 2b**). Besides showing better performance than the untreated mutant group, the mRNA-treatment of *GalT*-KO mice also demonstrated a return to the expected performance of WT animals (p=0.71). To validate these findings, R software was used to run a model assessing the effect of treatment and gender on composite score performance. This model confirmed that untreated *GalT*-KO animals performed significantly worse than the treated *GalT*-KO group (p = 0.00345), though this difference was found to be mostly driven by the untreated female animals (p = 0.0377) (**Supplemental Table 1c**). Post-HOC analysis confirmed the significant improvements in mRNA treated *GalT*-KO female animals compared to their untreated counterparts (p = 0.0344) (**Supplemental Table 1d**).

**Table 1b.** Descriptive Statistics from Composite Phenotype Scoring Assessments.

| Treatment Group | Treatment Age | Assessment No. | vars | n | Mean | SD | Median | Trimmed | MAD | Min | Max | Range | Skew | Kurtosis | SE |
| --- | --- | --- | --- | --- | --- | --- | --- | --- | --- | --- | --- | --- | --- | --- | --- |
| WT Control | 3 weeks | 1 | 1 | 8 | 1.19 | 0.651 | 1 | 1.19 | 0.556 | 0.5 | 2.5 | 2 | 0.79 | -0.693 | 0.23 |
| Untreated *GalT*-KO | 3 weeks | 1 | 1 | 8 | 2.44 | 1.35 | 2.12 | 2.44 | 0.927 | 0.5 | 5 | 4.5 | 0.492 | -0.797 | 0.479 |
| Treated *GalT*-KO | 3 weeks | 1 | 1 | 8 | 1.28 | 1.32 | 0.875 | 1.28 | 0.927 | 0.25 | 4.25 | 4 | 1.28 | 0.322 | 0.466 |
| WT Control | 6 weeks | 1 | 1 | 8 | 0.781 | 0.7 | 0.625 | 0.781 | 0.741 | 0 | 2 | 2 | 0.435 | -1.32 | 0.247 |
| Untreated *GalT*-KO | 6 weeks | 1 | 1 | 8 | 2.25 | 1.35 | 1.88 | 2.25 | 0.741 | 0.5 | 4.5 | 4 | 0.515 | -1.29 | 0.477 |
| Treated *GalT*-KO | 6 weeks | 1 | 1 | 8 | 2.16 | 2.11 | 1.5 | 2.16 | 0.927 | 0 | 6.75 | 6.75 | 1.13 | -0.0184 | 0.747 |
| WT Control | 3 weeks | 2 | 1 | 8 | 1.09 | 0.706 | 1 | 1.09 | 0.556 | 0.25 | 2.5 | 2.25 | 0.697 | -0.691 | 0.25 |
| Untreated *GalT*-KO | 3 weeks | 2 | 1 | 7 | 1.71 | 1.36 | 1.25 | 1.71 | 0.371 | 0.75 | 4.75 | 4 | 1.49 | 0.546 | 0.516 |
| Treated *GalT*-KO | 3 weeks | 2 | 1 | 8 | 1.47 | 0.881 | 1.12 | 1.47 | 0.741 | 0.25 | 3 | 2.75 | 0.382 | -1.27 | 0.311 |

Considering the Rotarod and Composite Phenotype Scoring Test results, we concede that mRNA-treatment of 3-week-old *GalT*-KO mice does improve their motor function.

### Scoring Assessments

#### Improvement of motor functions by a single dose of *GALT* mRNA-therapy did not sustain far beyond 3 weeks

To see if the positive changes resulting from *GALT* mRNA treatment are sustainable (question **2**), we re-tested these mice after nine weeks, or when they reached 23 weeks of age (**Fig. 1**). For Rotarod testing, untreated WT control mice have a median latency of 130 seconds, while untreated *GalT*-KO mice have a median latency of 95.5 seconds (**Table 1a**). We see 71.4% of the untreated mutant mice performing under the WT median threshold, demonstrating a significant genotypic difference in performance distribution (p=0.049). Treated *GalT*-KO mice have a median latency of 137.45 seconds, and they show significant difference in their distribution when compared to the untreated *GalT*-KO cohort (p=0.0227). While the untreated mutant mice mostly performed under the WT median latency, we found that only 37.5% of the mRNA-treated *GalT*-KO group fell below this threshold. No difference is noted between WT and mRNA-treated mutant group distribution (**Fig. 3a**). A generalized linear model with Gaussian distribution confirmed treatment type to have significant influence on latency. Untreated *GalT*-KO animals were estimated to perform 14.44 seconds worse than WT controls and 32.41 seconds worse than treated *GalT*-KO animals **(Supplemental Fig 2)**. This model also substantiated results from Chi-Squared analysis, showing that treated *GalT*-KO group still has significantly higher latencies than their untreated counterparts, 12 weeks following the end of their treatment (p=0.0111) (**Supplemental Table 2a**). Again, post-HOC testing in R confirms this distinction. Though a significantly better performance is still observed in the treated *GalT*-KO group at this time, the magnitude of effectis notably smaller than obtained in the Assessment 1. Comparing the two assessments, there is no apparent decline in the mRNA-treated *GalT*-KO group performance. Instead, we see a trend of increased latency in the untreated mutant group over time. With these results, we infer that the benefits from mRNA treatment of *GalT*-KO animals may still exist at this point, though the impact is surely decreasing.

**Figure 3.**
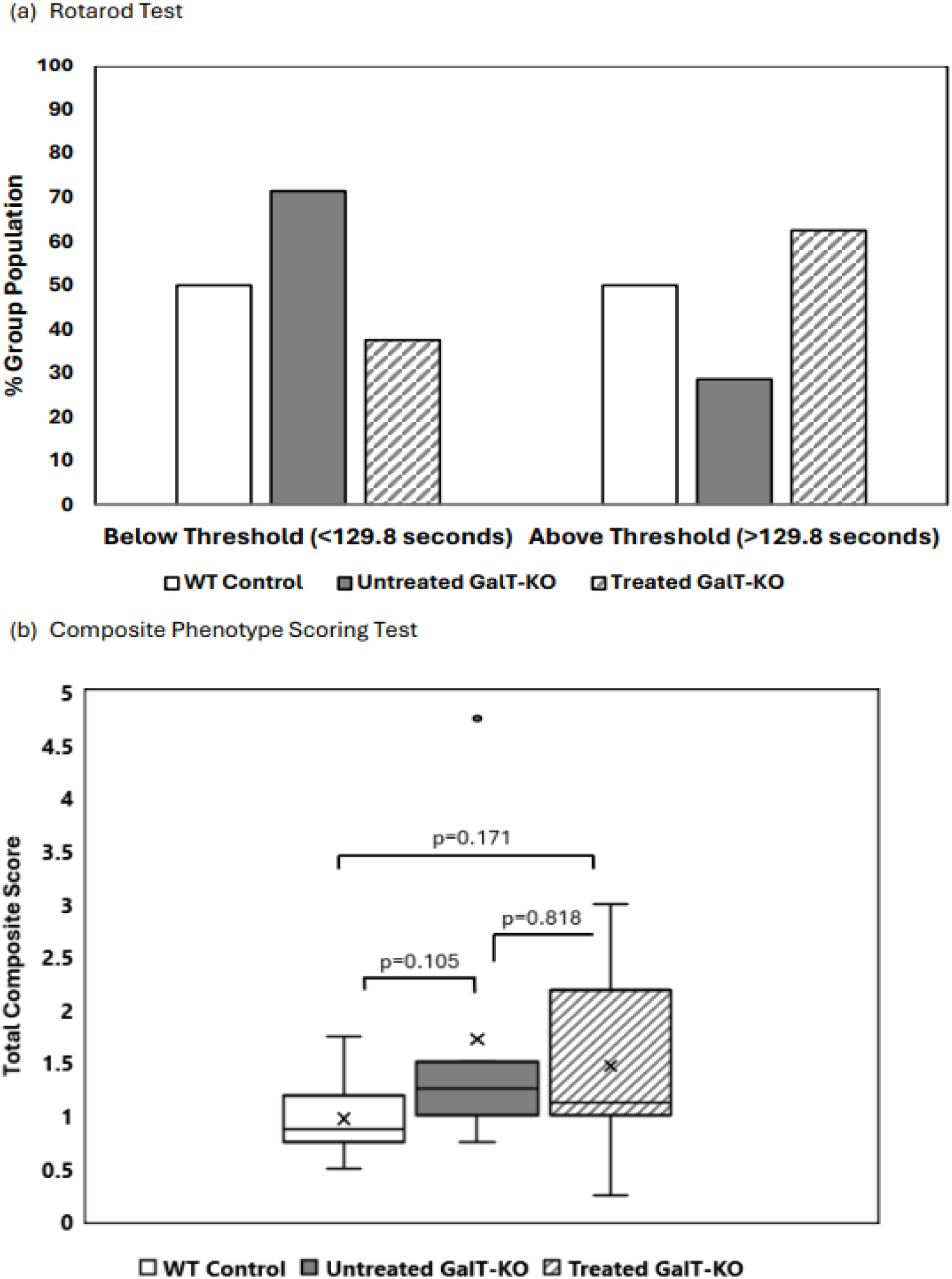
Motor function evaluation at 12 weeks after *GALT* mRNA treatment ended. (a) Rotarod performance distribution of WT, treated and untreated animals. The majority Treated *GalT*-KO animals performed above the WT control group median latency, whereas the untreated mutant animals mostly performed under this threshold. Chi-Squared Tests were performed to determine differences in latency distribution. (b) Composite Phenotype Scoring Test shows no significantly different performances between treatment groups. WT and untreated *GalT*-KO groups demonstrate an improved composite score from the first assessment, while mRNA-treated animals maintain their performance. Data represent mean ± SEM. Statistical significance was determined using Mann Whitney U test. General linearized models with gaussian distribution were run in R to determine the effect of treatment type on Rotarod and/or Composite Scoring test performances. (See Supplemental Data).

For Composite Score Phenotype Scoring Test, untreated WT mice demonstrated a score near to their first assessment, with a median total score of 0.875. Untreated and mRNA-treated *GalT*-KO groups performed similarly with median composite scores of 1.25 and 1.125, respectively (**Table 1b**). Compared to Assessment 1, both WT and untreated *GalT*-KO groups improved their scores, while the treated mutant group showed slight decline in performance (**Fig 3b**). There were no distinctions between any of the three groups in this assessment, as confirmed with the Mann Whitney U test and models run in R (**Supplemental Table 2c**). Consequently, we cannot confidently confirm the benefits of mRNA treatment in *GalT*-KO animals have sustained over time for this Test.

### *GALT* mRNA-treatment appears to be more effective in early life

Finally, we compared the above results with separate groups of mice that we began dosing at an older age to address Question 3 (i.e., 6 weeks of age) (**Fig. 1**). For Rotarod testing, untreated WT control mice have a median latency of 126.85 seconds, while untreated *GalT*-KO mice have a median latency of 101.85 seconds (**Table 1a**). Despite the mRNA-treated *GalT*-KO group showing a slightly higher median latency (122.55 seconds) than their untreated counterparts, both treated and untreated *GalT*-KO groups follow the same performance distribution relative to the WT threshold latency. We cannot differentiate these groups as 66.7% of each population performed under the WT median latency (**Fig. 4a**). A generalized linear model confirms the lack of influence mRNA-treatment has on latency in this assessment (**Supplemental Table 3a**). While the treatment of 6-week-old *GalT*-KO animals did not show any improvements in rotarod performance, we see that the untreated *GalT*-KO animals performed much better in the 6-week versus 3-week cohort.

**Figure 4.**
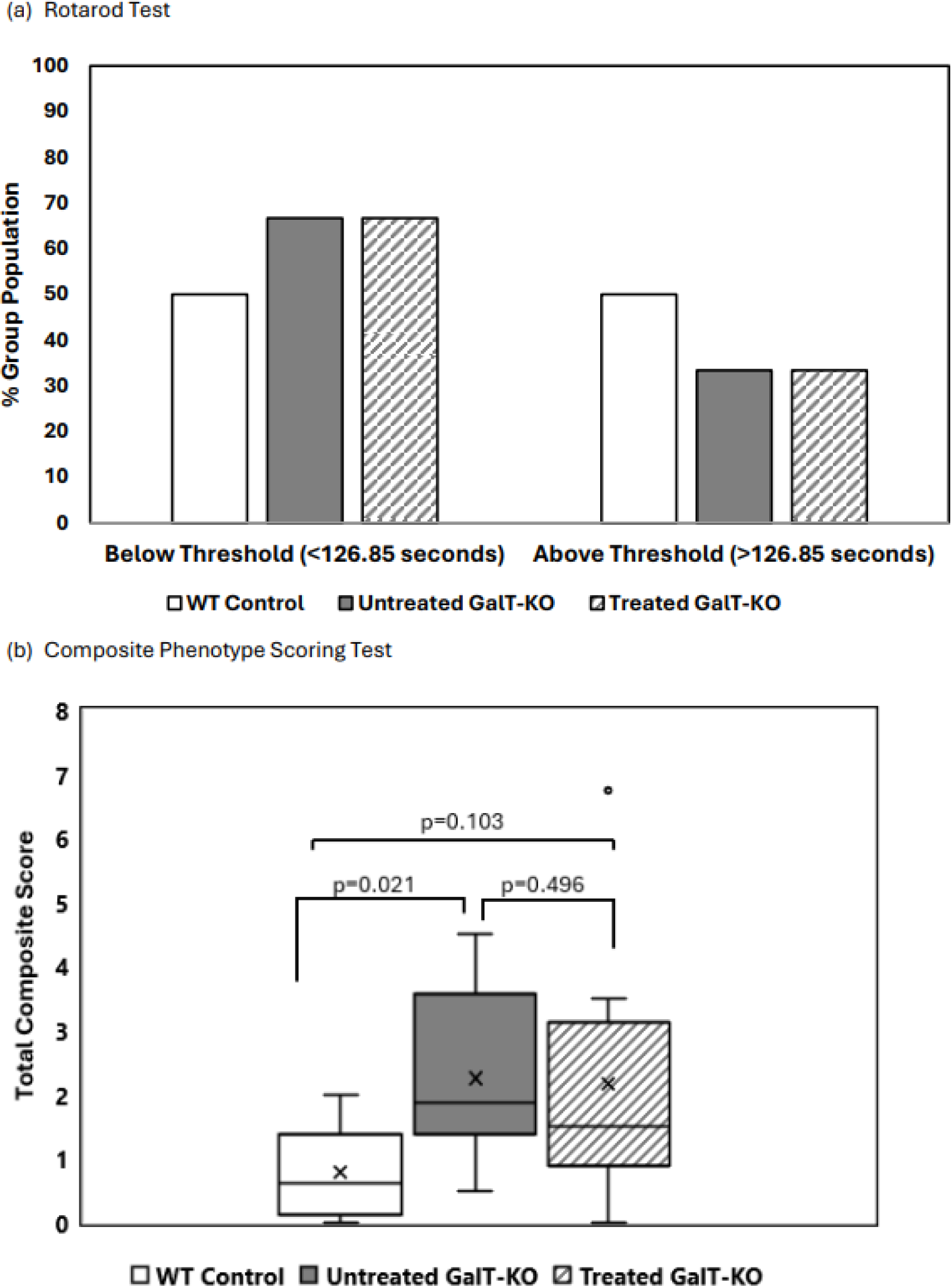
Motor function assessment in aged mice following GALT mRNA treatment administered at 6 weeks of age. (a) Rotarod test results demonstrate no difference between treated and untreated *GalT*-KO performance relative to WT median latency. Chi-Squared analysis was utilized to determine significance in rotarod performance distribution. (b) Composite Phenotype Score evaluation indicates there is no comparable performance between untreated *GalT*-KO mice and mRNA-treated *GalT*-KO animals. Data represent mean ± SEM. Statistical significance was determined using Mann-Whitney U test.

For the Composite Phenotype Scoring test, untreated WT control mice have a median total score of 0.625, and untreated *GalT*-KO mice have a significantly higher median score of 1.875 (p=0.021) (**Fig. 4b**). mRNA-treated *GalT*-KO mice performed similarly, with a median total score of 1.5 (**Table 1b**). Unlike the untreated mutant group, however, the treated *GalT*-KO animals show no significant deviation from WT median scores (**Fig. 4b**). Neither treatment group nor gender have a significant influence on composite scoring abilities, as confirmed by generalized linear models run in R (**Supplemental Table 3c)**.

## 4. Discussion

Recent advances and successes in molecular therapeutics have paved the way for revolutionary treatment modalities for many monogenic recessive diseases like Phenylketonuria and Classic Galactosemia. Among these modalities, mRNA-based therapies are gaining ground because the successful track record in vaccine development. Additionally, such treatment, if successful, will address the root cause of the diseases, thus making them rational choices when one aims for a potentially curative therapy. In this study, we aimed to extend our earlier *in vivo* POC work for an experimental *GALT* mRNA therapy for Classic Galactosemia from disease-relevant biomarkers ^17^/ functional biomarker ^18^ normalization to disease-relevant phenotypes correction in a mouse model of Galactosemia. As we have previously shown that these *GalT*-KO animals manifest motor-related impairment,^22^ we focused on the potential improvement of this phenotype.

As a pilot investigation, we chose to test one dosing regimen, which is a biweekly dosing of 2mg/kg for two months before we commenced the behavioral assessments (Rotarod and Composite Phenotype Scoring tests). We chose bi-weekly administration because it will affect the compliance of patients if they have to receive such treatment more than once in every two weeks. We selected 2mg/kg because it was determined in our earlier studies that such dosage is safe and a single IV dose of this specific *GALT* mRNA is effective in maintaining a normal level of galactose metabolites in liver of the mutant mice up to 2 weeks.[18] We opted for a two-month-long treatment because we wanted a short-term multi-dose study. Enrolled animals were dosed starting at 21- and 42-days old *via* tail-vein injections, repeating biweekly for two months. Behavioral assessments were conducted at 3- and 9-weeks following the final mRNA dose to assess motor impairment phenotypes. Baseline behavioral assessments were not carried out for this study because the animals will be too young, and many potentially confounding physiological changes will occur after the dosing is completed. Instead, results were directly compared between age- and sex-matched treatment groups. Moreover, previous experiments conducted by our lab suggest that motor-impairment phenotypes as assessed by Rotarod and Composite Phenotype Scoring tests are not detectable until our mouse model reaches at least breeding age (6-8 weeks). This was another reason for us proceeding without pre-treatment assessments.

We were able to see some significant improvements in Rotarod and Composite Phenotype Scoring tests in the first assessment of the younger mutant mice treated with the *GALT* mRNA, when they were 14 weeks of age (**Fig. 2**). Assessment 1 demonstrated a clear distinction between untreated WT and *GalT*-KO animals, and a return to normal behavior in the mRNA-treated *GalT*-KO group. By applying these results to Question 1, we determined that *GALT* mRNA therapy *does* improve motor-related phenotypes in our mouse model. Several modes of data analyses were used to assess results from both behavioral tests, in which all showed significant improvement in the mRNA-treated *GalT*-KO animals compared to untreated mutant mice.

Paradoxically, we saw an improvement in the performances of the untreated mutant mice for both Rotarod Test (i.e., increased latency) and Composite Phenotype Scoring Test (i.e., lower combined score) at Assessment 2 when they reached 23 weeks of age (close to 6 months) (**Fig. 3**). Although such improvement has wiped out much of the genotypic differences between WT and untreated *GalT*-KO mice that we saw in the first assessment (**Fig. 2**), the mRNA treated animals continued to perform above the WT median latency and significantly outperformed the untreated *GalT-*KO animals (p=0.0227) (**Fig 3a**). Comparing the Rotarod Assessments 1 and 2, we do not detect any change in performance in the mRNA-treated animals, indicating that the treatment effects have likely sustained. Although there is no notable decline in performance in the treated group, we do see the gap between untreated and treated *GalT*-KO animals is much smaller. In Assessment 1, we observed extremely high significance when testing the difference between treated and untreated *GalT*-KO groups (p=0.000179), and untreated *GalT*-KO animals were estimated to perform 49.37 seconds worse than treated animals (**Supplemental Table 1a**). In the Assessment 2, we still see a substantially worse performance in the untreated *GalT*-KO animals compared to treated *GalT*-KO, though to a much smaller degree with the untreated group estimated to score 32.41 seconds lower (p = 0.0111) (**Supplemental Table 2a**). mRNA-treatment of *GalT*-KO animals seems to sustain its impact on Rotarod test and allows treated mutant mice to perform at WT levels nine weeks after treatment was finished. Though the improvement is still visible, we see a declining trend where the difference between treated and untreated animals is becoming less overtime.

In the Composite Phenotype Scoring Assessment 2, the improved scores initially seen in the mRNA treated animals were no longer detectable (**Fig 2b**). We also puzzlingly noted superior scores in the untreated mutant group over time. (**Fig.3b)**. e. However, the data together suggested that the positive changes we witnessed in the first assessment began to wane, which is unsurprising due to the finite half-life of mRNA.

To answer question 2, although some improvements from mRNA treatment of *GalT*-KO animals have sustained over this period in Rotarod testing, the Composite Phenotype Scoring test results suggest that the beneficial effects of treatment have since worn off. It would be interesting to do more regular and prolonged testing on another group of animals following this dosing regimen, to determine how long we can expect mRNA treatment to sustain its benefits to motor-related phenotypes.

Results from **Fig. 4** revealed that for the dosing regimen tested, it is more effective to commence the mRNA treatment at a younger age (i.e., 3-week-old). Similar to what we found in **Fig. 3**; the older untreated *GalT*-KO mice tested at 17 weeks old appeared to perform quite well at Rotarod test. The mRNA-treated mutant animals did not show any significant improvement over the untreated mutants. Instead, we see an identical distribution of “under-performing” and “over-performing” mice in the untreated and mRNA-treated *GalT*-KO mouse groups (**Fig. 4a**). For Composite Phenotype Scoring Test, we did see a genotypic difference between the untreated *GalT*-KO and the WT mice (p=0.021). Like the Rotarod Test, however, *GalT*-KO mice treated at 6 weeks of age do not appear to have any improvement for the Composite Score test (**Fig. 4b**). No significant influence of treatment was detected when running linear regression models on these results (**Supplemental Table 3b**).

Despite being a small-scale study, our data indicated that when treated early in life, the experimental *GALT* mRNA is effective in improving the motor-related phenotypes in the *GalT*-KO mice using the specified dosing regimen. In addition, our data suggested that due to the finite half-life of mRNA, repeated dosing should be considered to sustain the long-term positive changes. We concede that we have not performed any correlation with the various biochemical biomarkers and GALT activity/protein abundance in these animals. However, we have done extensive pharmacokinetic and pharmacodynamic analyses of our specific *GALT* mRNA species in the previous work,^17, 18^ and therefore, we focused on the phenotypic analyses in this study. Finally, it would be interesting to extend the current study to other disease-relevant phenotypes like subfertility in the future.

## Author Contributions

Study concept and design: KL, BB, OB, and AG. Animal study data collection, data analysis, data graphing: OB, AJ BB and KL. Statistical analysis: OB and AJ. Manuscript writing: KL, BB, OB, JS, and XY

## Data Availability Statement

Data supporting the studies presented in this manuscript can be made available by request to the corresponding authors.

## Acknowledgments

K.L. is supported by the Eunice Kennedy Shriver National Institute of Child Health and Human Development grants (R01HD089933, R21HD104056).

## Conflicts of Interest

The authors declare the following competing interests: X.Y., and J.S., are employees of Moderna and hold equities from the company.

