## Supplemental Figures for "Experimental galactose-1 phosphate uridylyltransferase (GALT) mRNA therapy improves motor-related phenotypes in a mouse model of Classic Galactosemia"

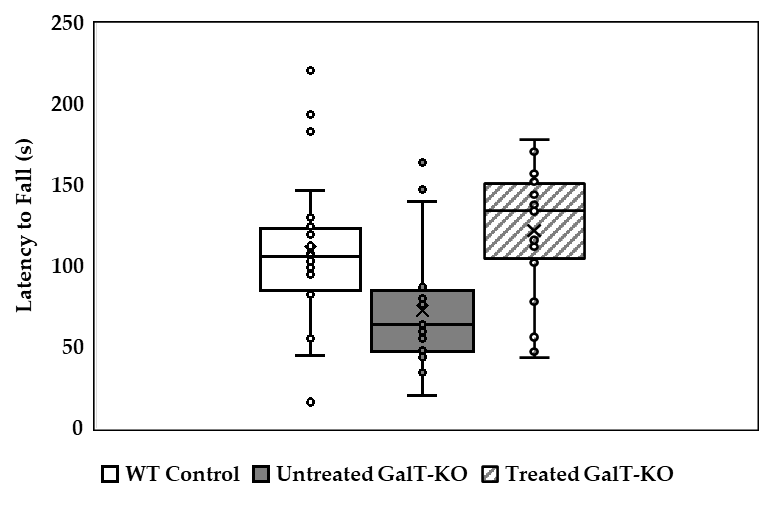
**Supplemental Figure 1.** Latency distribution of 3-week treatment groups.

**Fig 1.** Boxplot showing the latency distribution for Cohort 1. *GalT*-KO animals treated at 3 weeks with mRNA display the highest latency out of the three groups, followed by WT controls. Untreated *GalT*-KO animals performed the worst in Assessment 1.

**Supplemental** **Table 1.**

(a)
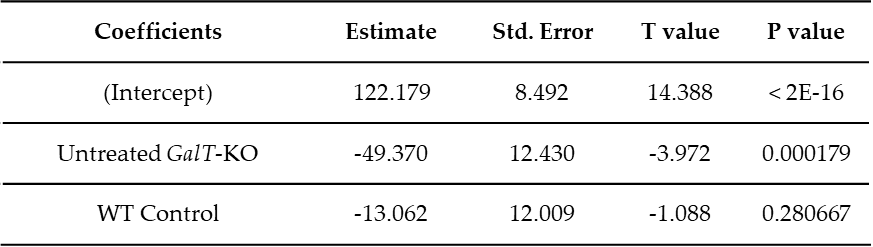
Generalized Linear Model (GLM) with Gaussian distribution assessing the effect of treatment type on latency, using mRNA-treated *GalT*-KO animals as the reference group
(b) Post-HOC analysis on 3-week Rotarod model findings.
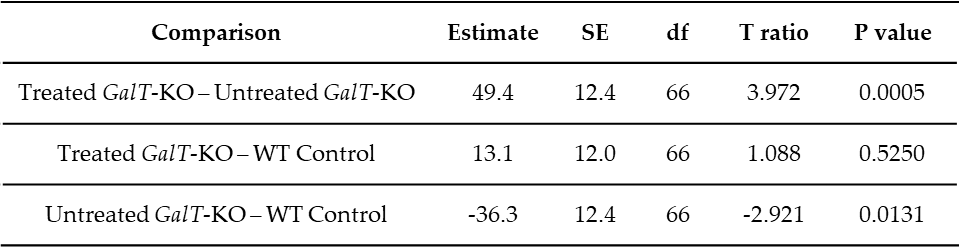

(c)
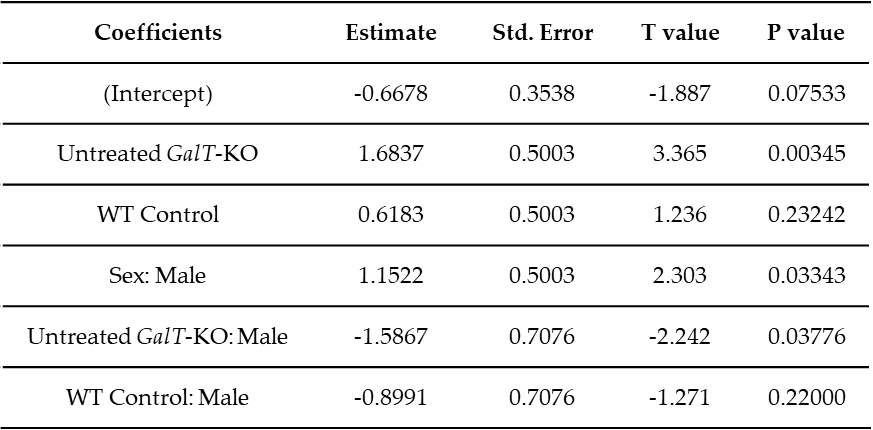
 3-week Composite Phenotype Scoring Test General Linearized Model (Box-Cox transformed data).
(d)
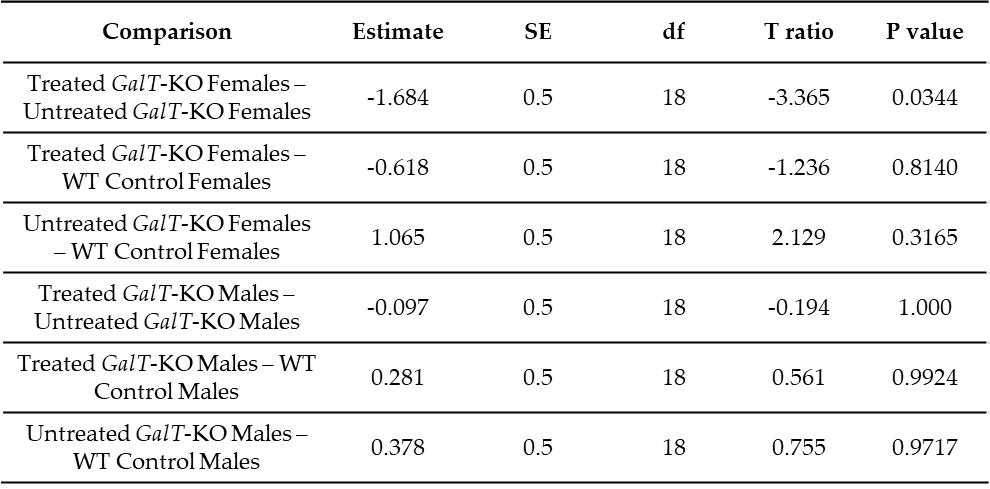
Post-HOC analysis on 3-week Composite Phenotype Scoring Test GLM findings.

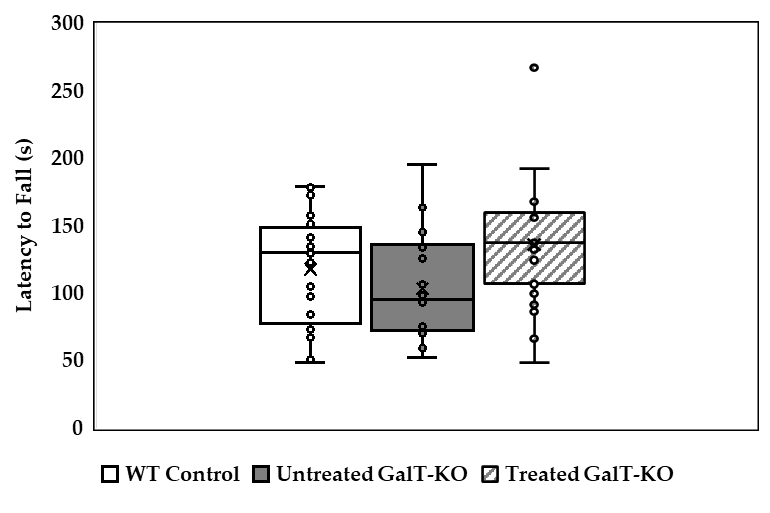
**Supplemental Figure 2: Latency distribution by treatment group in Assessment 2.**

**Fig 2.** Treated *GalT*-KO mice demonstrated significantly longer latency times compared to untreated mutant animals (p = 0.0111).

**Supplemental Table 2.**

(a)
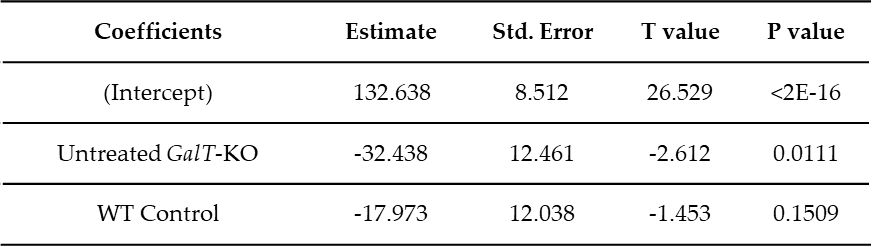
 GLM with Gaussian distribution assessing Rotarod performances in Assessment 2.
(b)
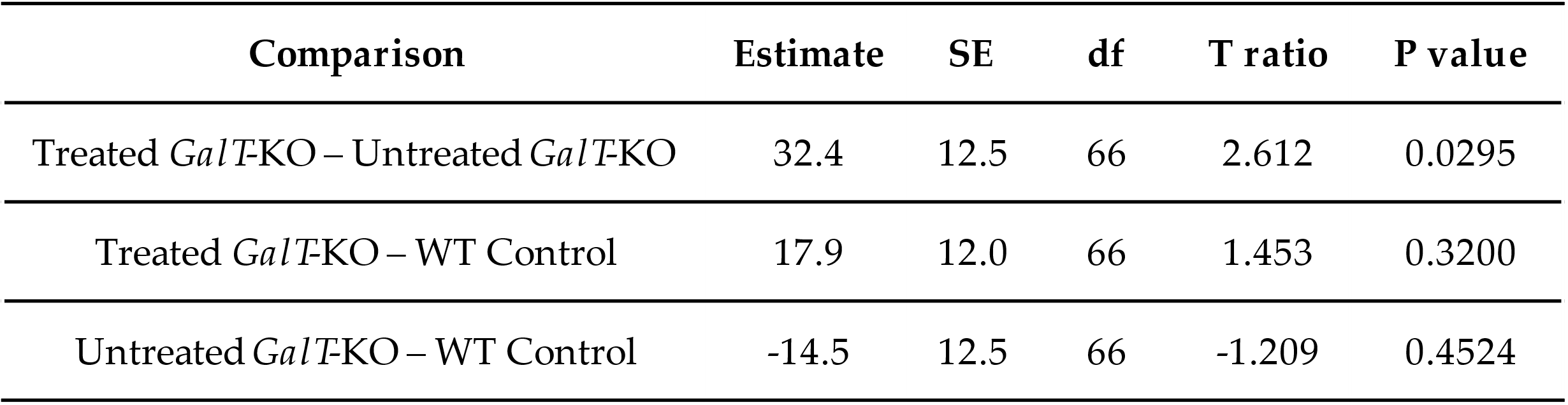
Post-HOC analysis on Assessment 2 Rotarod GLM.
(c)
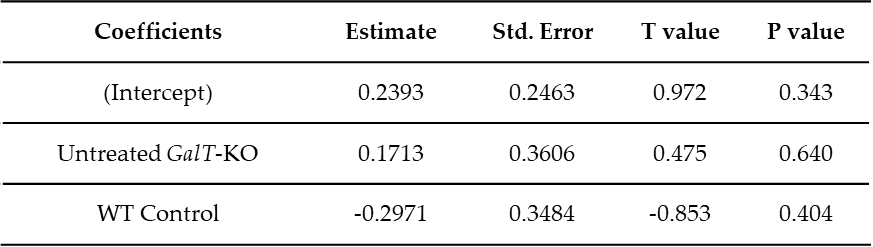
 GLM assessing Composite Phenotype Scoring Test results and Treatment interactions in Assessment 2.

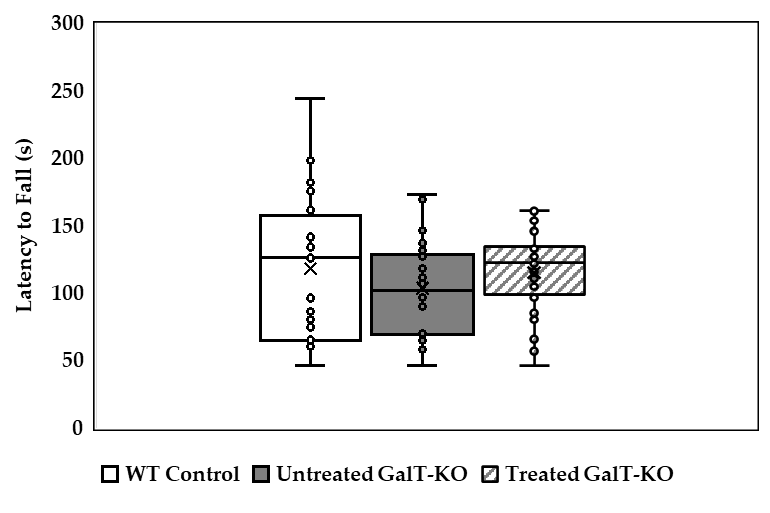
**Supplemental Figure 3. Rotarod performance of 6-week treated animals.**

**Supplemental Table 3.**

(a)
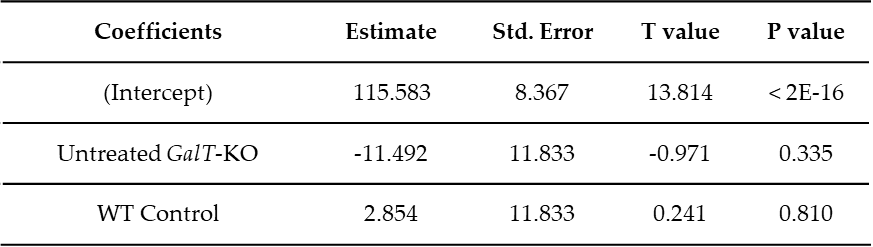
6-week group Rotarod Generalized Linear Model results.
(b)
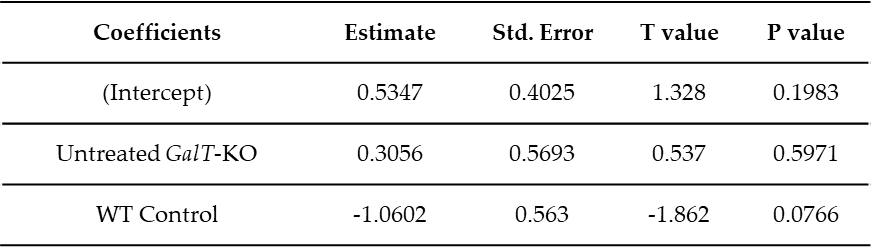
GLM results for Composite Phenotype Scoring Test of 6-week treated animals.
